# Contrasting dynamics and biotic association networks in estuarine microbenthic communities along an environmental disturbance gradient

**DOI:** 10.1101/2025.01.07.631661

**Authors:** Leire Garate, Anthony A. Chariton, Ion L. Abad-Recio, Andrew Bissett, Anders Lanzén

## Abstract

Estuarine ecosystems experience a range of anthropogenic pressures. Consequently, robust monitoring tools are essential for their management and protection. Utilising environmental DNA in routine monitoring programs enables the inclusion of benthic microorganisms, which are not only good indicators of environmental condition, but also play critical roles in ecosystem functioning. In this study we collect eDNA from sediment samples across time in six estuaries, from the Basque coast (Spain), under varying degrees of disturbance. To identify potential indicators of biotic integrity (environmental health status), we used time series data to examine the prokaryotic microbial communities and consensus networks associated with different levels of pollution. In general, sediment communities were relatively temporally stable, with the moderately and heavily disturbed sites showing more variation. The consensus networks also differed significantly in their topologies, with more impacted estuaries having fewer nodes, edges and connectance, among others, and higher modularity compared to those less impacted. Moreover, the potential keystone taxa and predicted functional profiles differed between consensus networks. This illustrates how modelled association networks can reveal new insights regarding the state of estuarine ecosystems and their potential functional processes.

## INTRODUCTION

Known as *itsasadar* (“sea branch”) in the Basque language spoken in our study area, estuaries are where the sea enters a riverbed and its surrounding marshlands and intertidal areas with the tide. This makes them dynamic, complex and productive ecosystems where marine, freshwater and specific estuarine biomes meet (Kennish 2002; Suzzi et al. 2023b). Estuaries provide valuable ecosystem services such as coastal protection, nurseries fish species, cross-boundary subsidies and removal of pollutants (Pinto and Marques 2015; Raes et al. 2022; Suzzi et al. 2023a). However, the sharp increase of human population and exploitation of coastal areas have severely impacted estuaries worldwide, through eutrophication, industrial pollution and hydrological changes, reducing some to little but tidal canals (Barbier EB et al. 2011; Boerema and Meire 2017).

Environmental monitoring is fundamental for providing information on the state of estuaries and other coastal habitats, thus informing management, protection and conservation of these areas. Biological components of ecosystems, in particular, represent and respond to impacts and other environmental parameters, integratively and cohesively, thus reflecting past and present environmental conditions. However, current approaches typically suffer from high costs and poor scalability, as well as a limited understanding of estuarine ecology, particularly of the microbial communities, which are central to the functioning of these ecosystems, in concert driving important processes such as degradation of pollutants and nutrient cycling (Lanzén et al. 2020; DiBattista et al. 2024; Moreira et al. 2024). Due to their sensitivities to a range of stressors, small size, functional versatility and ubiquity within sediments, benthic prokaryotic microorganisms are excellent indicators of ecosystem health (Aylagas et al. 2017; Birrer et al. 2019; Cordier 2020; Moreira et al. 2023).

Utilising eDNA derived data, Lanzén et al. (2020) previously identified a number of bacterial taxa as potential indicators for a range of different types of anthropogenic pollutants in sediments. These indicators were developed based on samples from the routine monitoring of Basque estuaries and resulted in the development of preliminary biotic indices as well as machine learning models predictive of environmental impact and ecological quality (EQ, indicating ecosystem health). However, the ecological roles played by most of these taxa remain unclear. One way to uncover the potential roles of taxa within the microbial communities in which they occur, is to model ecological association networks based on co-variance of individual taxon abundances over temporal or spatial replicates. However, this approach is prone to generating taxon associations (or “co-occurrences”) resulting from shared responses to spatial or seasonal variation in environmental variables rather than direct interactions (Faust et al. 2015). In any case, using time series to infer association networks has the advantage of capturing the dynamics and seasonality of the studied community, in which taxa are both responding to and recovering from perturbances in a correlated manner. Modelling also suggests that this approach is more likely to result in correct inference of biological interactions (Fisher and Mehta 2014).

Association networks are valuable tools for understanding the potential interactions and importance of individual taxa in a community, which can be valuable for understanding their ecological roles (Lima-mendez et al. 2015; Garate et al. 2022). Equally important, network topology can reveal properties of the community as a whole, and act as a multifaceted bioindicator in itself. For example, ecological networks from communities under stress typically show a higher modularity (Veloso et al. 2023) and share of positive to negative associations (Hernandez et al. 2021). These observed connections between topological properties and EQ are not surprising, considering the fundamental role of both competitive as well as mutualistic and syntrophic interactions for community composition (Zelezniak et al. 2015).

Here, we aim to identify potential indicators of estuarine EQ by studying the sediment prokaryotic communities from six estuaries from the Basque coast. These communities were assessed in terms of their composition, temporal stability and associations between taxa. Estuaries were sampled twice monthly for at least 12 months. Apparent seasonal trends in these microbenthic communities were relatively weak or absent, especially compared to coastal plankton communities from nearby sites (Garate et al. 2022). This allowed us to better model the potential biological interactions and contrast them between different communities. We did this by choosing sites with contrasting levels of environmental impact, pairwise merging networks from sites with similar EQ into *consensus networks.* Although some studies have addressed seasonality in estuarine microbenthos to a limited extent (Böer et al. 2009; Zhang et al. 2020b), few, if any have monitored community structure over time in a seasonal timescale and utilised the resulting data for inference of ecological association networks.

## METHODS

### Estuarine sediment sampling

Six routinely monitored intertidal sites of the Basque Monitoring Network (BMN; Borja et al. 2016) were selected for this study (Figure 1). Sites were selected so that they formed three pairs, each sharing similar ecological status, known from previous monitoring based on macroinvertebrates using AMBI (Borja, Franco and Pérez 2000) and M-AMBI (Muxika, Borja and Bald 2007), 16S rRNA gene metabarcoding of prokaryotic microbenthos (Lanzén et al. 2020, Table 1) and physicochemical parameters.

**Fig. 1.**
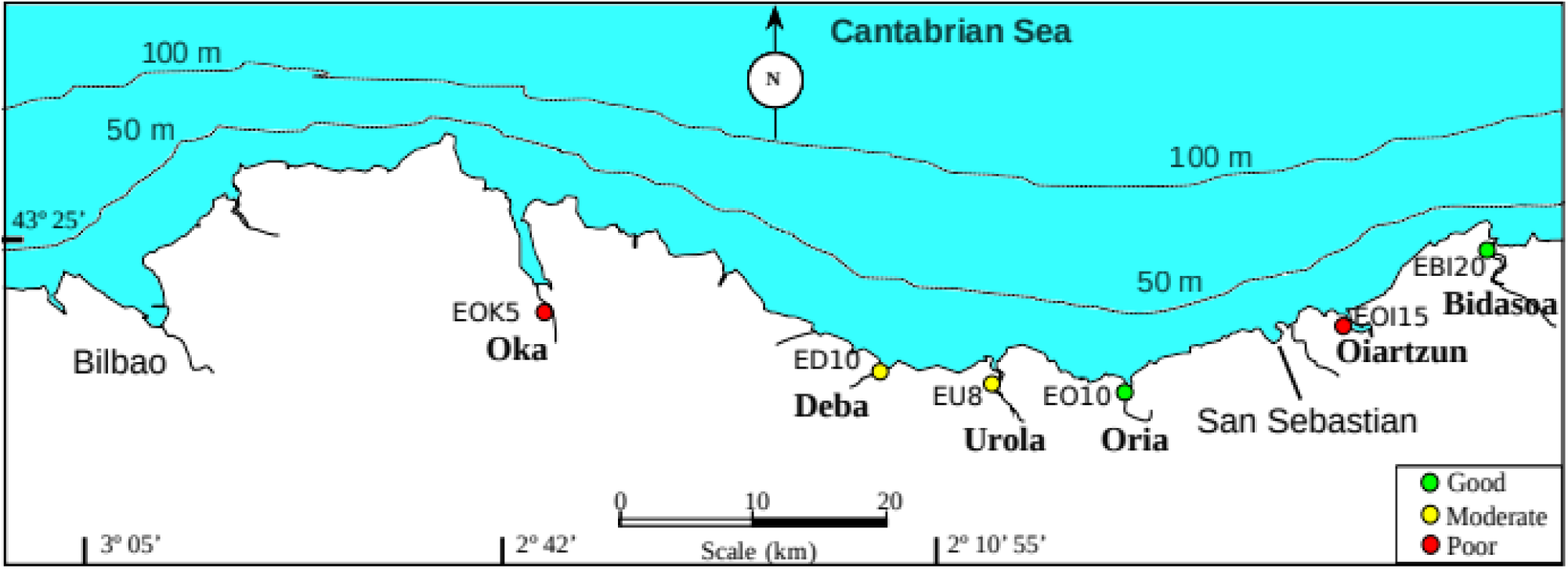
Map of the Bay of Biscay where the six estuaries are located. Colours represent the ecological qality (EQ) of the estuaries.

**Table 1.** Description of the study sites of the time series with number of time points (TPs), in order of ecological quality (EQ)

| Site | Estuary | #TPs | EQ | Main pressures | Characteristics | Location: UTMX, UTM Y (ETSR89) |
| --- | --- | --- | --- | --- | --- | --- |
| EBI20 | Bidasoa | 30 | Good | Urban, treated wastewater | Mesohaline, sandy | Hondarribia: 598024, 4802583 |
| EO10 | Oria | 31 | Good | Urban, treated wastewater, industry | Polyhaline, sandy | Orio zahar Beach: 570456, 4792570 |
| ED10 | Deba | 22 | Moderate | Urban, treated wastewater, industry | Mesohaline, shifting (gravel — mud) | 1866 Deba bridge: 552146, 4793495 |
| EU08 | Urola | 31 | Moderate | Treated wastewater, industry (incl. upstream papermill) | Mesohaline, muddy, oxygen depleted | Zumaia industrial park: 561251, 4793515 |
| EOI15 | Oiartzun | 22 | Poor | Commercial harbour, wastewater from food processing, industry | Polyhaline, muddy, oxygen depleted | AZTI Pasaia, Trintxerpe: 586667, 4797168 |
| EOK05 | Oka | 22 | Poor | Untreated (primarily only) urban wastewater (until 2022) | Oligohaline, muddy, oxygen depleted | Downstream Gernika WWTP: 527059, 4798683 |

From each site, sediment samples were collected twice monthly starting mid-2019 to early 2020, until late 2020 or early 2021, with a minimum of 23 time points and one full year coverage per site (Table 1). Samples were collected by scraping the top centimetre of the sediment into a sterile 50 ml Falcon tube and were kept on ice in a portable cooler until arrival to the laboratory. All the samples were kept at −80 °C until nucleic acid extraction.

### Nucleic acid extraction and metabarcoding

Total DNA was extracted using either RNeasy PowerSoil DNA Elution Kit (Qiagen) or DNeasy PowerSoil Pro Kit, following the manufacturer’s instructions. Two different kits were used due to the interruption on the production and availability of RNeasy PowerSoil kits during the COVID-19 pandemic. No significant differences were observed in terms of yield and composition between kits (Supplementary Figure 1). Nucleic acid extracts were visualised on an electrophoresis gel and their concentration and quality were measured with a Nanodrop spectrophotometer and Qubit dsDNA BR Assay Kit, used in a Qubit fluorometer. Sample concentrations were normalised to 10 ng/µl for 16S rRNA amplification.

The V4 region of the 16S small subunit rRNA gene was amplified with primers 519F (CAGCMGCCGCGGTAA; Øvreås et al. 1997) and 806RD (GGACTACNVGGGTWTCTAAT; Apprill et al. 2015) as described previously by Lanzén *et al*. (2020). Two negative controls per plate were added before DNA extraction and PCR. Triplicates of each sample were amplified as follows: 95 °C for 3 minutes, 30 cycles of 95 °C for 30 s, 50 °C for 30 s and 72 °C for 45 s, and a final extension at 72 °C for 5 minutes. PCR triplicates were then pooled and purified using AMPure XP beads (Beckman Coulter). 5 µl of purified amplicons were used as template for a second PCR round in which Illumina Nextera adapters were attached to the amplicons. The program used in the second PCR was: 95 °C for 3 minutes, 8 cycles of 95 °C for 30 s, 55 °C for 30 s, and 72 °C for 30 s, followed by a final extension at 72 °C for 5 min. The second PCR product was again purified using AMPure XP beads and eDNA concentration was measured using a Qubit fluorometer. PCR products yielding sufficient amount and quality of amplified DNA were then pooled in into a final library at equimolar concentrations of 10 nM. Products yielding insufficient quantities for this pooling (negative controls) were added columetrically 5 µl. Sequencing was carried out using Illumina MiSeq with paired-end 2×300 bp v3 chemistry, carried out at the National Genome Analysis Centre (CNAG - Centre Nacional d’Anàlisi Genòmica, Barcelona) and distributed over four different runs. Raw sequences are available from the INSDC Sequence Read Archive with BioProject accession number PRJEB83451.

### Sequence analyses

Sequence data processing was performed as described previously by Garate et al. (2022). Briefly, read-pair overlapping and sequence data filtering was carried out using vsearch v2.7.1 (Rognes et al. 2016) allowing 20 mismatches, and primers removed using cutadapt v1.18 (Martin 2011), followed by truncation to 252 bp, discarding reads without full and correct primer sequences or more than one expected error. The remaining sequences were de-replicated and sorted by abundance using *vsearch* and clustered into sequence variants (SVs) using *SWARM* v2.21 (Mahé et al. 2015). Abundances of unique sequences across samples were retained to construct an SV contingency table, using the scripts *fasta_merging.py* and *matrix_creation.py*, of *SLIM* (Dufresne et al. 2019). We then discarded SVs with a total abundance across samples of one (singletons), then putative chimeras, using *vsearch*, reference based with SilvaModPR2 v138 (Lanzén et al. 2012); https://github.com/lanzen/CREST) followed by de novo mode. To correct remaining PCR and sequencing artefacts and to merge intra-specific or intra-genomic SVs, we then applied LULU post-clustering curation (Frøslev et al. 2017) with default parameters except for increasing minimum similarity to 97%). Curated SVs were aligned to SilvaModPR2 v138 using *blastn* v2.6.0+ and then taxonomically classified using *CREST* v3.1.0 (Lanzén et al. 2012).

Using R, SVs unclassified at kingdom rank and non-target SVs originating from Eukaryota (18S or organellar SSU; n=5651) were then removed along with 47 putative (p<0.05) contaminant SVs, identified based on PCR and extraction blanks, with *decontam* v.1.12.0 (Davis et al. 2018). Cross-contamination was reduced by setting OTU abundances to zero where SVs occurred in a sample at an abundance being 100x lower relative to its average non-zero abundance across samples (corresponding to the UNCROSS algorithm, Edgar 2016). To decrease bias from uneven sequencing depth across samples, all rare SVs defined as those that never reached 0.1% abundance in at least one sample, were removed. Finally, technical replicates resulting from comparisons of extraction kits or repeated DNA extractions were pooled *in silico,* leaving 158 time-point samples across the six sites (see Supplementary Table 1).

Filtered SVs with identical taxonomic annotation (and their abundances across time-point samples) were merged into operational taxonomic units (OTUs) based on taxonomy, for reconstruction of association networks. This was done by merging all SVs with identical annotations, without a fixed taxonomic rank, meaning that our final list includes both species rank OTUs (e.g. *Desulfuromusa bakii*) as well as “orphan” OTUs for the parent genus or higher ranks summarising all SVs unclassified at species or lower ranks (e.g., *Arenimonas*, Sedimenticolaceae.).

All used metadata, final SV and OTU distribution tables, and scripts that are not part of a publicly available free software packages, are available from http://github.com/lanzen/EstuarySedimentTS.

### Consensus association network reconstruction and visualisation

Ecological association networks for each site were based on relative abundances of taxonomy-based OTUs present in at least 50% of the time points of that site. Networks were calculated with *wTO* v. 1.6.3 (Gysi et al. 2018) for R (R Core Team 2021) as described in Garate et al. (2022). The resulting site-specific networks belonging to the same ecological status group were then used to calculate consensus networks (CNs) for Good (EBI20+EO10), Moderate (EU08+ED10) and Poor (EOK05+EOI15) EQ groups (hereafter Good, Moderate and Poor CN, respectively). The function *wTO.Consensus* of the *wTO* R package used to calculate the CN only retains associations which appear in all the overlapped networks and that have the same sign. Only correlations with wTO*_CN_* > 0.4 and estimated p<0.001 were retained, with the resulting CNs exported to *Cytoscape* v.3.9.1 (Shannon et al. 2003).

Networks were analysed using the Cytoscape tool Analyse Network, using the R packages *igraph* (Csardi and Nepusz 2006) and *NetIndices* (Kones et al. 2009). Potential keystone taxa were defined as previously described by Garate *et al*. (2022), selecting the top ten nodes with the highest sums of their degree of connectivity (number of connections to different nodes) and closeness centrality (how close a node is to all others). The ten nodes with the highest betweenness centrality were defined as connectors. Modules were identified using *ClusterMaker2* in Cytoscape (Morris et al. 2011) based on the Glay community method.

### Network comparison

The online tool *Netshift* (Kuntal et al. 2019) was used to compare consensus networks. This software identifies ‘drivers’ as the nodes that gain importance when the ecosystem is perturbed. This importance is based on a NESH score calculated by the program and the increment in the betweenness centrality of such nodes. In essence, if a node has a sufficiently high NESH score and its scaled betweenness centrality increases from the Control to the Case networks, then it is considered a ‘driver’. Thus, we identified driver taxa using: (i) the Good CN as control with the Moderate CN as case; and (ii) the Moderate CN as control with the Poor CN as case, to identifying drivers to a worse EQ. We also used (iii) the Poor CN as control with Moderate as case; and (iv) Moderate CN as control with Good as case, to identify drivers that lead the changes to an improved EQ. The drivers were selected as those taxa with NESH score > 2 and an increase in the scaled betweenness centrality.

### Potential indicator taxa identification

We identified potential indicator taxa as those taxa that both were keystones (play important roles in the ecosystems) and drivers (important when the ecosystems are changing its ecological status).

### Metabolic potential prediction

PiCRUSt2 (Langille et al. 2013; Douglas et al. 2020) was used to infer the potential functional roles of the bacterial communities from the six study sites. In our case, two analyses were carried out: one general to obtain a list of normalized abundances of the potential KOs present in the community, and a second one to ascertain the contribution of each taxon of that abundance in each of the sample. PICRUSt2 (v.2.5.2) was used with default settings to place phylogenetically taxa and infer their function according to KEGG Orthology (KO). Placed sequences with Nearest Sequenced Taxon Index (NSTI) scores > 2 were removed. The --*strat_out* flag was also used, to identify the specific taxa contribution to each of the predicted KOs. The resulting table containing the normalized abundance of each KO and e taxon that contributed the most to that abundance was filtered in order to study the metabolic potential of the nine taxa that were identified both as keystones in the CN and drivers to the same ecological status. SIMPER tests were carried out on these results to look for KOs that caused differences in the microbial communities.

### Statistical analyses

Data was analysed and represented using *vegan* (Oksanen 2018), *phyloseq* (McMurdie and Holmes 2013), *microbiome* (Lahti and Shetty 2017), and *ggplot2* (Wickham 2011) packages in R. The Shannon alpha diversity index was calculated using the alpha function of the *vegan* package in R, Chao1 richness was calculated with the richness function of the *microbiome* R package. Pairwise Wilcoxon tests were used to check significant differences in diversity and richness among estuarine ecological states. Pairwise Bray Curtis dissimilarities were calculated with the *vegdist* function. NMDS plots (function metaMDS) based on Bray Curtis dissimilarity were performed to compare estuarine microbiomes with different ecological states, after Hellinger transformation of the data. Differences between microbial communities from the three disturbance levels were analysed using the ANOSIM function from *vegan* package. SIMPER analyses were calculated on one hand to identify the taxa responsible of such dissimilarities between microbiomes, and on the other hand, to identify the KOs whose normalized abundance differed among the three categories of disturbance.

## RESULTS

### Microbial composition and diversity

Across the six study sites, we obtained approximately 9.8 million sequence reads after filtering. The two sites with lower disturbance and Good EQ (EBI20 and EO10) appeared to have the most taxonomically stable microbial communities over the time series length, dominated by Gammaproteobacteria (∼50% of relative abundance), Bacteroidota (∼ 25% of relative abundance) and Alphaproteobacteria (∼ 15%) (Figure 2). Thaumarchaeota and Actinobacteroidota were also relatively abundant in these two sites (∼ 10% each), as well as in the Moderate EQ site ED10. In EBI20 (Good EQ), Thaumarchaeota appeared to show some seasonality, with lower relative abundances during the summer. Gammaproteobacteria was also the most relatively abundant class in the least and moderately disturbed sites (EU08 and ED10, 50% and 40-50%, respectively), with a sharp increase of Desulfobacterota in ED10 in the summer months (up to 40% of relative abundance), which appeared to coincide with the occurrence of a thicker overlying muddy top layer above the otherwise sand and gravel dominated riverbed. Desulfobacterota was more relatively abundant at all sites regardless of EQ, being highest at EOI15 and least in EBI20. In EOI15, Campylobacterota (25-40 %) was equally relatively abundant to Gammaproteobacteria and Desulfobacterota (25 % each), while in the other Poor EQ site, EOK05, Campylobacterota only became abundant during the last winter months (Figure 2).

**Fig. 2.**
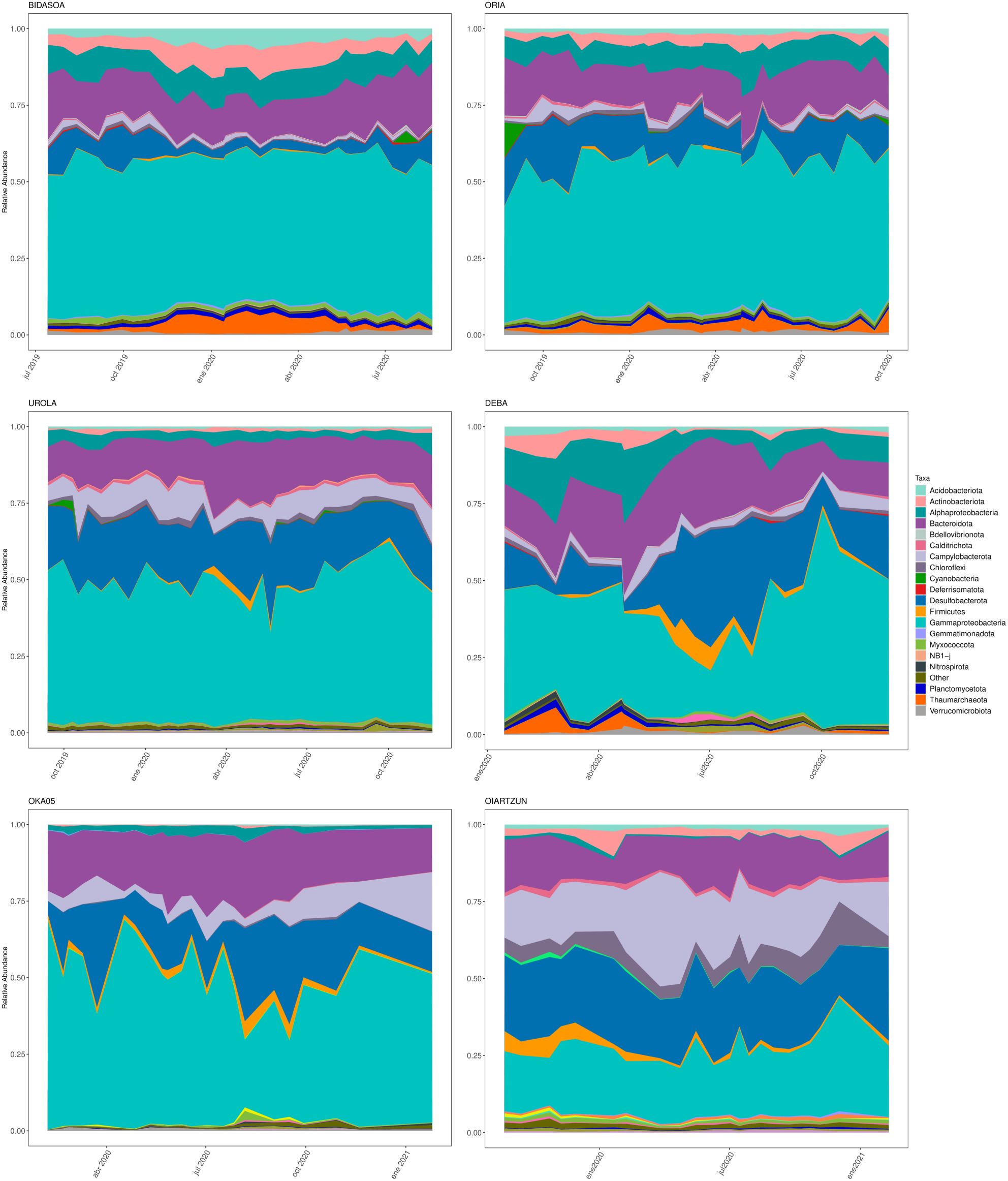
Relative abundance of 20 most abundant phyla across the temporal series in each of the estuaries. The top two correspond to the good EQ estuaries, the middle two correspond to the intermediate EQ estuaries and the bottom two to the poor EQ estuaries.

Shannon diversity index and Chao1 richness were significantly lower in heavily disturbed sites, (Supplementary Figure 1). Pairwise comparisons (Wilcoxon test) indicated significant differences between the Shannon diversity of the three EQ groups (p<0.001), although the Chao1 was significantly different only between the heavily disturbed compared to the two other categories (p<0.001).

An NMDS representation of community dissimilarities in each site throughout the time series confirmed the grouping by EQ, with very clear separation between Poor and Good EQ sites, while the moderate EQ site ED10 was more variable and overlapped with the distributions of both Good and Poor sites (Figure 3A). ANOSIM analysis confirmed the significant differences between the three types of sites (Good EQ-Intermeditate EQ R=0.624, p= 0.001; Good EQ-Poor EQ R=0.835, p=0.001; Intermediate EQ-Poor EQ R=0.826, p=0.001). SIMPER analyses identified 175 taxonomic OTUs that significantly differed in relative abundance between these three groups, including 15 phyla. Among these, Acidobacteriota, Actinobacteriota, and Thaumarchaeota appeared to decrease with higher disturbance, while Chloroflexi, Campylobacterota, Desulfobacterota and Firmicutes increased (Supplementary Figure 2).

**Fig. 3.**
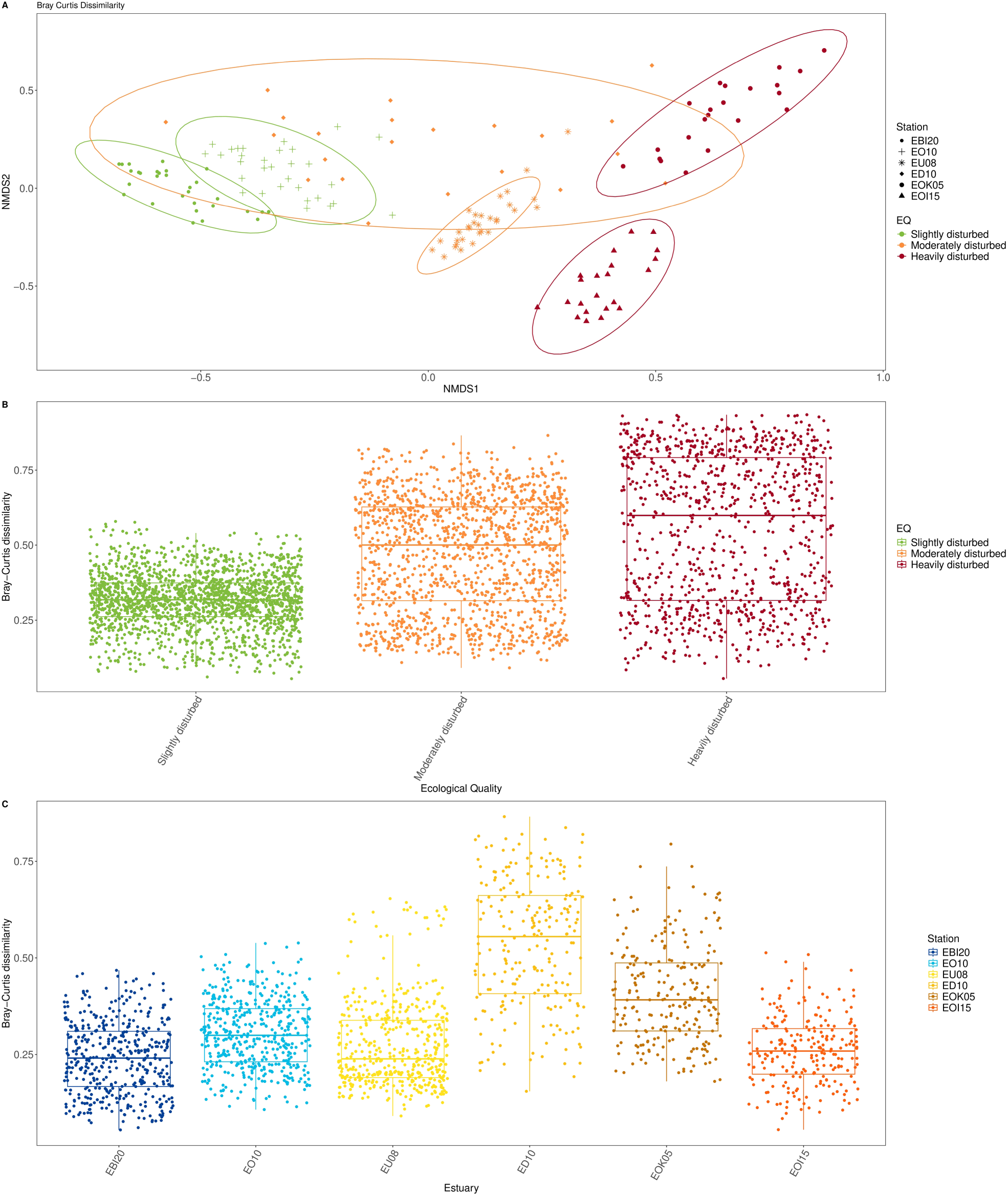
A) NMDS representation based on Bray Curtis dissimilarities for the prokaryotic communities in the six estuaries colured by EQ. B) Pairwise dissimilarities between time points for the estuaries grouped by EQ. C) Pairwise dissimilarities between time points for individual estuaries.

The distribution of pairwise dissimilarities between time point samples from sites of the same EQ indicated increased variability with higher disturbance, i.e., worse EQ (Figure 3B). Due to including two sites per EQ, this measure includes both temporal variability in the same site as well as differences between the two sites. Restricting the analysis to only time points from the same site revealed a less clear trend with the moderately disturbed site ED10 having the highest temporal variability, followed by the heavily disturbed EOI15 (Figure 3C).

### Network analyses and identification of key taxa

Consensus networks (CNs) for the three EQ groups (Figure 4) showed clear differences in terms of several topological properties (Table 2). The Good CN was more complex, denser and centralised with more nodes and edges, as well as a higher connectance, clustering coefficient, centralisation and maximum degree, while the Poor CN had the lowest connectance, centralisation, maximum degree, and numbers of nodes and edges. The share of negative edges was also highest in the Good CN and lowest in the Poor, while modularity showed the opposite trend, increasing with disturbance. The Moderate CN consistently showed intermediate values for these properties, more resembling those of Good than Poor status.

**Fig. 4.**
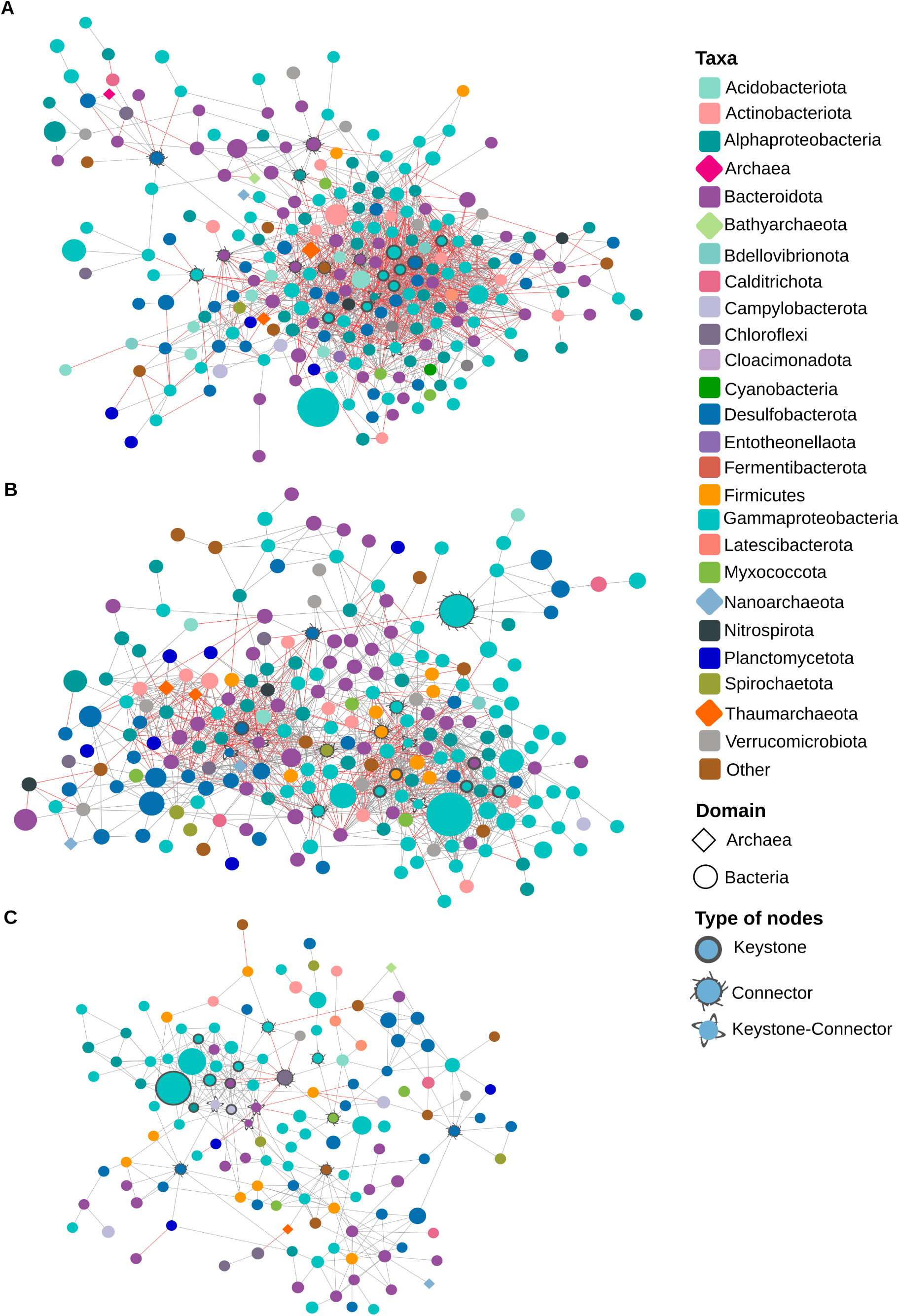
Modelled consensus networks for those estuaries with the same EQ. A) Good EQ estuaries, B) Intermediate EQ estuaries, C) Poor EQ estuaries. Nodes represents taxa and their size correspond to the mean relative abundance across the time series. Nodes are coloured by phyla. Gray lines connecting nodes represent positive associations and red lines represent negative associations.

**Table 2.** Properties of the EQ group consensus association networks.

| Property / EQ | Good | Moderate | Poor |
| --- | --- | --- | --- |
| #Nodes | 243 | 230 | 135 |
| #Edges | 1433 | 1195 | 340 |
| Negative edges | 523 (36%) | 278 (23%) | 21 (6%) |
| Connectance (network density) | 2.4% | 2.3% | 1.8% |
| Modularity | 33% | 50% | 59% |
| Maximum degree | 52 | 39 | 18 |
| Centralisation | 17% | 13% | 10% |
| Heterogeneity | 1.03 | 0.84 | 0.84 |
| Clustering coefficient | 0.35 | 0.39 | 0.34 |

Keystones and connector nodes (Table 3) generally showed low mean abundances, except for a unique keystone in the Poor CN (Rhodocyclaceae). In the Good CN, nine keystones belonged to Proteobacteria phylum and one to Desulfobacterota, while the keystones of the Moderate and Poor CNs were more taxonomically diverse, belonging to four and three different phyla, respectively (see Table 3). Some of the keystones were also connectors, meaning that they play an additional role connecting modules within the network, namely two in the Good CN (Hyphomonadaceae and *Halioglobus),* four in the Moderate CN (Steroidobacteraceae, Prolixibacteraceae, *Arenimonas* and *Desulfobacter*) and two in the Poor CN (Macellibacteroides and *Arcobacter*).

**Table 3.**
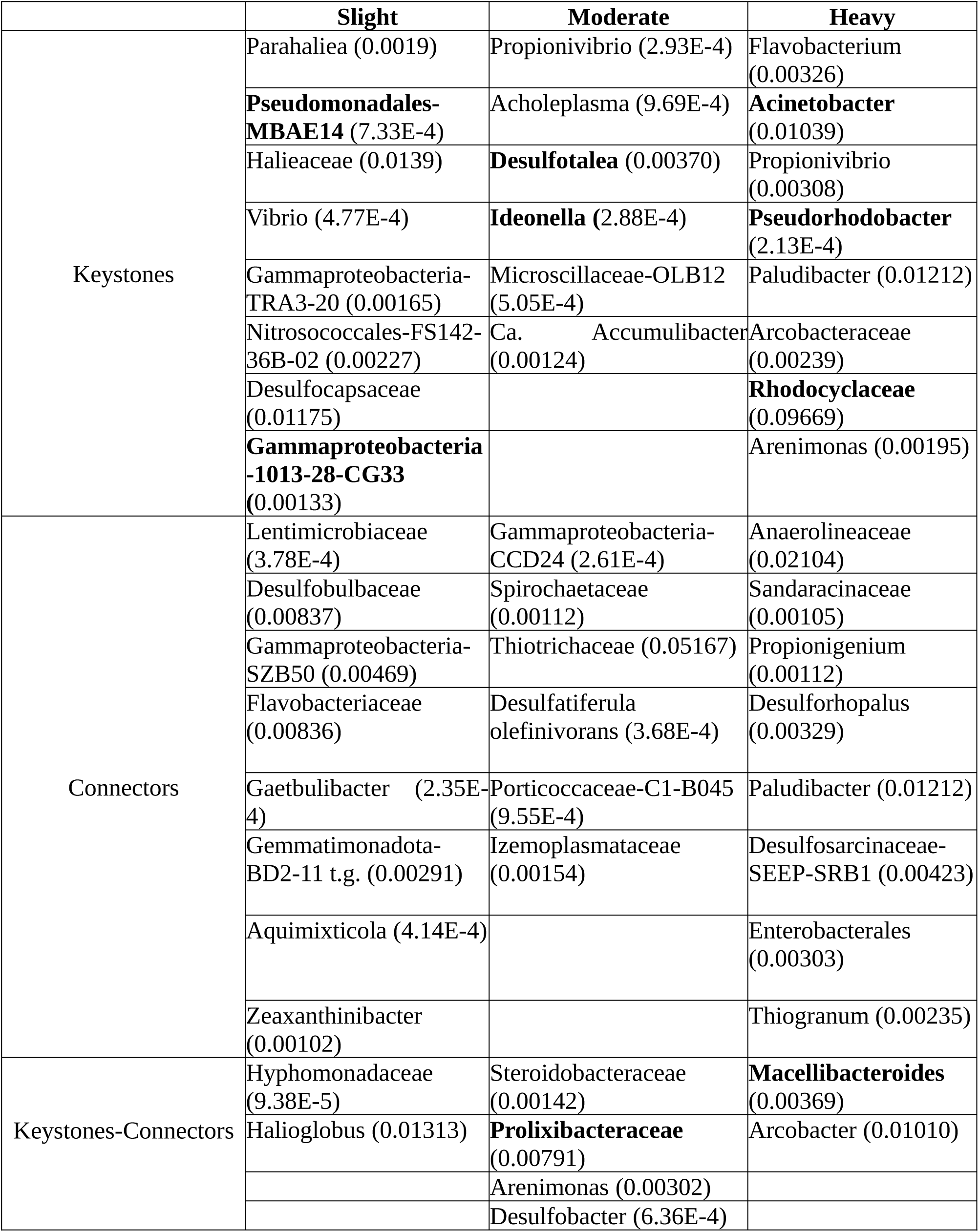
Taxa that are keystones, connectors, and keystones&connectors. Keystones are the top ten taxa that had the highest values of degree of connectivity plus clustering coefficient and connectors the top ten taxa with the highest betweenness centrality. In brackets it is shown the relative mean abundance of each taxa across the time series. Taxa in bold are those identified as potential indicators (keystones and drivers, see Table 4).

The four NetShift analyses performed (Figure 5) identifying the most significant “drivers” from one EQ to an adjacent one (Good to Moderate, Moderate to Bad, Bad to Moderate and Moderate to Good), resulted in different selections depending on the target EQ (Table 4). Those drivers that overlapped with keystones from the target CN were selected as potential indicator taxa, namely Pseudomonadales Clade MBAE14 and Gammaproteobacteria Clade 1013-28-CG33 (Intermediate to Good); *Ideonella* (Poor to Intermediate), *Desulfotalea* and Prolixibacteraceae (Good to Intermediate); and *Pseudorhodobacter*, *Acinetobacter*, Rhodocyclaceae and Macellibacteroides (Intermediate to Poor).

**Fig. 5.**
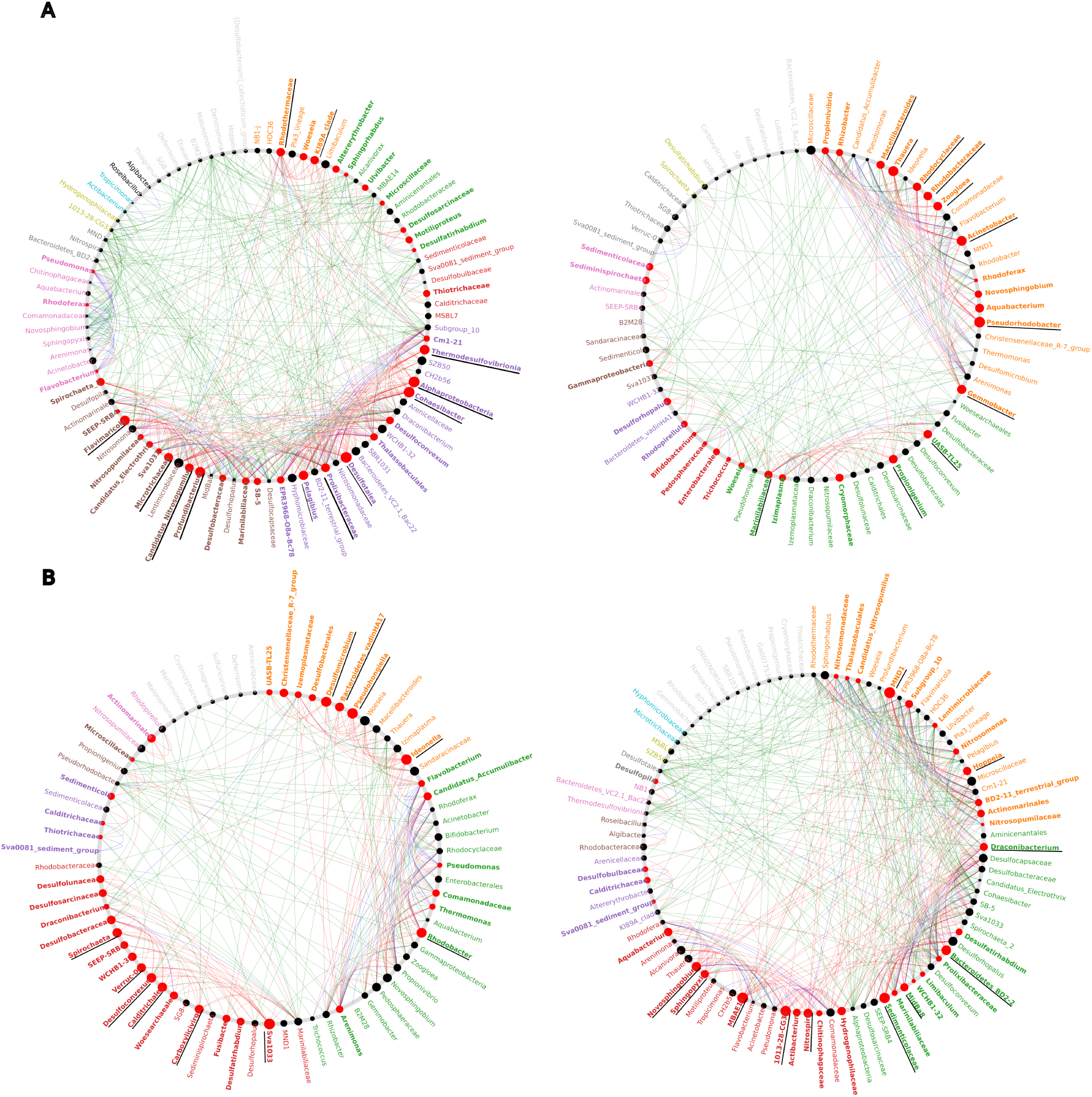
Netshif analyses. A) The left panel shows the result of comparing the good EQ (control) and Intermediate EQ (case); the right panel shows the result of comparing the Intermediate EQ network (control) with the Poor EQ network (case). B) the left panel shows the comparison between the Poor EQ (control) with the Intermediate EQ (case); right panel shows the comparison between the Intermediate EQ (control) with the Good EQ (case).

**Table 4.** Drivers identified as those taxa with NESH score >2 and increase of the scaled betweenness centrality. Taxa marked in bold are taxa identified as potential indicators, as they are keystones and drivers.

| Good to Intermediate | Intermediate to Poor | Poor to Intermediate | Intermediate to Good |
| --- | --- | --- | --- |
| Alphaproteobacteria | Thauera | Desulfuromonadales-Sva1033 | Nitrosomonadaceae-MND1 |
| Cohaesibacter | Gemmobacter | Pseudohongiella | Sedimenticolaceae |
| Profundibacterium | Propionigenium | Desulfomicrobium | Bacteroidetes BD2-2 |
| Flavimaricola | Zoogloea | Rhodobacter | Novosphingobium |
| Thermodesulfovibrionia | Rhodobacteraceae | Desulfoconvexum | Nitrospira |
| Pelagibius | Marinilabiliaceae | Calditrichales | Hoppeia |
| Microtrichaceae | Polyangia-UASB-TL25 | Spirochaeta 2 | Draconibacterium |
| Ca. Nitrosopumilus | **Pseudorhodobacter** | Bacteroidetes vadinHA17 | Polyangia-MidBa8 |
| Rhodothermaceae | **Acinetobacter** | Puniceicoccaceae-Verruc-01 | Sphingopyxis |
| Pseudomonadales-KI89A clade | **Rhodocyclaceae** | Carboxylicivirga | Hydrogenophilaceae |
| **Desulfotalea** | **Macellibacteroides** | **Ideonella** | **Pseudomonadales-MBAE14** |
| **Prolixibacteraceae** |  |  | **Gammaproteobacteria-1013-28-CG33** |

### Potential metabolic activity

The results of PiCRUSt2 analyses for analysing the potential KOs, without considering taxa contribution, did not retrieve significant differences between the estuarine microbial communities. However, when studying the data from the taxa contributions focused on keystone-driver taxa, we obtained a total of five KOs (KO0830, KO1322, KO1270, KO0116, KO1077) that had statistically significant differences among groups according to the disturbance level (slightly, moderately and heavily disturbed). The highest normalized abundances of these KOs occurred in the sites corresponding to the EQ of the keystone-driver taxa that contributed most to each KO (Figure 6).

**Fig. 6.**
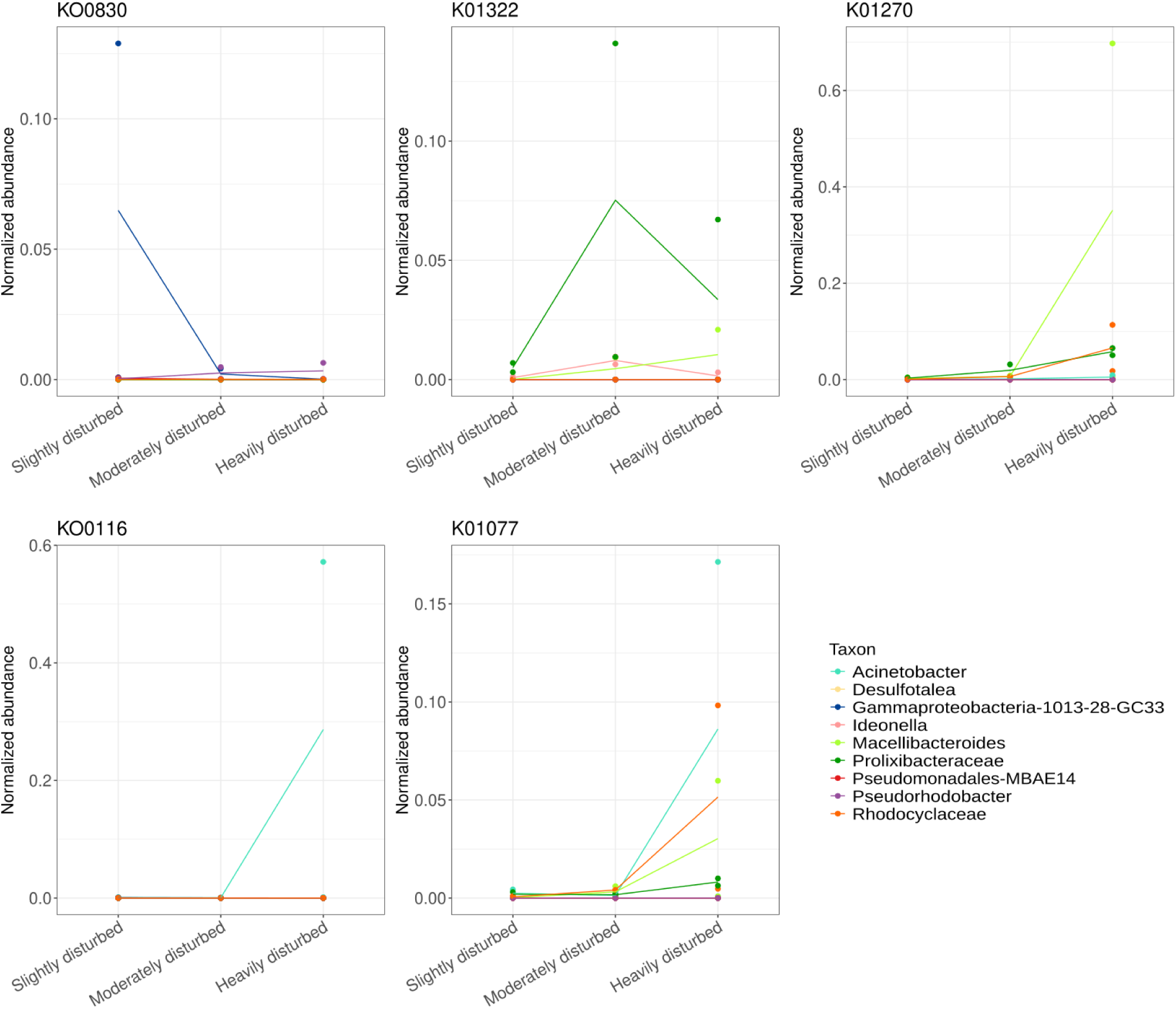
Normalized abundance of the five KOs with significant differences when considering the contribution of the nine potential indicators taxa identified.

## DISCUSSION

In this study, we monitored the microbenthic communities of six estuarine sites under contrasting degrees of impact from coastal pollution and eutrophication (two sites with Poor EQ, two with Moderate and two with Good) twice monthly for little over a year, comparing their dynamics and modelled association networks. We show that less impacted sites in general were more stable over time, and dynamics did not appear to be seasonal. Network topologies, predicted keystone, connector and “driver” taxa also differed clearly, especially in the Poor EQ communities. Finally, we identified potential indicator taxa, of interest for biological monitoring, predicted to drive the change in interaction structure as well as acting as keystones.

### Estuarine microbiota: diversities and composition

The Poor EQ sites presented lower values of alpha diversity Shannon Index and extrapolated richness according to the Chao1 index, in agreement with several other studies in marine sediments and soil microbiomes (Böer et al. 2009; Kim et al. 2013; Hestetun, Lanzén and Dahlgren 2021; Zhang et al. 2023), although other studies have shown the opposite results, where the diversity increased under low levels of physical disturbance (Galand et al. 2016), or higher metal concentration (Salah-Tantawy et al. 2022). It is worth noting that Shannon diversity in our study was higher in the Moderate EQ sites, as observed in several studies and often framed, although ambiguously and likely incorrectly, as the Intermediate Disturbance Hypothesis (Fox 2013). In ED10, sediment properties varied strongly during the time series, shifting from a sandy to a thin mud dominated top sediment during spring and early summer, above a gravel dominated riverbed. The top sediment also varied in thickness and sometimes appeared completely absent in large areas. This was clearly reflected in the higher temporal beta diversity (variability) in this site and in the microbial composition and agrees with previous studies regarding how different sediment types structure prokaryotic communities (Boey et al. 2022; Zvi-Kedem et al. 2024).

Spatial variability of marine meio- and macrobenthos has also been suggested as an indicator of negative environmental impact (Warwick and Clarke 1993), in agreement with the combined effects of temporal and spatial variability observed in this study for microbenthos. However, limiting the analysis to individual sites makes the intermediate EQ site ED10 an outlier due to its higher temporal variance, while the heavily disturbed site EOI15 hosted a relatively stable microbenthic community temporally. Thus, a larger dataset incorporating time series from more sites would be necessary to better evaluate the correlation between variability and impact.

Taxonomically, all six sites except the highly disturbed site in Oiartzun (EOI15) were dominated by different taxa belonging to the Gammaproteobacteria. Especially, unclassified Gammaproteobacteria, Thiotrichaceae and *Woeseia,* in less and moderately disturbed sites (including *W. oceani* in EBI20). Woeseiaceae are known to be widespread and abundant worldwide in marine sediments ranging from the deep sea to the coast. They have wide functional versatility, ranging from facultative sulfur- and hydrogen-based chemoautotrophy to obligate heterotrophy (Mußmann et al. 2017; Buongiorno et al. 2020; Hoffmann et al. 2020). Actinobacteriota, Thaumarchaeota and Planctomycetota also appeared to be more sensitive to disturbance, appearing only in the less disturbed sites, but also in the Deba estuary (ED10). This agrees with several previous studies. For example, Thaumarchaeota, chemoautotrophic ammonia oxidizers of central importance to nitrogen cycling in freshwater and marine systems, have previously been found to be enriched in nutrient limited sediments (Marshall et al. 2021) and sensitive to pollutants such as crude oil (Urakawa et al. 2019), mixed industrial pollution and eutrophication including in the same estuaries as studied here (Lanzén et al. 2020).

As expected, based on previous studies, Desulfobacterota, Campylobacterota and Firmicutes tended to be more abundant with higher disturbance. Desulfobacterota, specifically Desulfocapsaceae and *Desulfosarcina* have previously been identified as indicators of mercury contamination (Rincón-Tomás et al. 2024), polluted mudflats (Ye et al. 2023), nutrient enrichment in salmon aquaculture sites (Keeley, Wood and Pochon 2018) and of poor EQ in the same estuaries as studied here, especially in sediments with negative redox potential (Lanzén et al. 2020).

The communities of the two most contaminated sites differed more from each other compared to other sites from the same EQ, in spite of having similar status as predicted by AMBI and M-AMBI (Borja et al. 2021). However, Oiartzun, in addition to nutrient enrichment, is subjected to significant levels of PAHs and Hg, with detectable levels of PCBs and DDTs, suggesting that the ecosystem impact or disturbance to the microbial community may be underestimated by the macrofauna-based biotic indices. Further, the impact on the studied site is likely more local due to its proximity to a greywater outlet, offering connectivity to less impacted sediments in a way that the Oka site does not. The difference may also be explained by their contrasting salinities, with the site in Oiartzun being polyhaline and that in Oka being oligohaline. The former showed higher abundances of sulphur oxidising Campylobacterota, and Chloroflexi (mainly Anaerolineaceae), compared to other sites. Chloroflexi are known to be involved in the degradation of hydrocarbons (Kragelund et al. 2007) which may explain their presence in the Oiartzun site. They have previously been encountered in high abundance in a disturbed coal sludge-receiving site (Zhang et al. 2023) and nutrient enriched sediments (Ye et al. 2023), and in particular Anaerolineaceae identified previously as an indicator of environmental impact in Basque estuaries (Lanzén *et al*. 2020), along with the dominant genera of Campylobacterota in this study (*Sulfurimonas* and *Sulfurovum*). Presence of simultaneous dominance of both Campylobacterota and Desulfobacterota suggests an intra-benthic sulphur cycle, with both dissimilatory sulphate reduction and oxidation of sulphide taking place within the sampled top sediments.

### Ecological networks analyses, comparison and identification of key taxa

Ecological consensus networks (CNs) showed differences in complexity, topology, keystones (highest degree centrality plus closeness centrality), connectors (highest betweenness centrality) and drivers (NESH score and increase in betweenness centrality), depending on EQ. In general, taxa deemed to be important to CN structures (keystones, connectors and divers), were generally of low relative abundance, in agreement with previous studies where the keystone taxa identified (based on number of associations) had relative abundances below 1% (Zhang et al. 2020a; Zhou et al. 2021). Further, we could not observe a correlation between taxon relative abundance and degree or centrality, suggesting that the risk is low for dominant taxa being spuriously over-represented among taxa inferred to be important for network structure, as suggested by Berry and Widder (2014) based on simulated data from a time series of a community with known interactions.

Several topological properties of CNs showed trends that agree well with previous studies based on association networks. For example, higher modularity with ecosystem stress has previously been reported (Veloso et al. 2023) and has been suggested to contribute to the stability of ecosystems subjected to environmental impacts such a s pollution, limiting severe impacts to specific modules and thus preventing community collapse (Stouffer and Bascompte 2011). Veloso et al. (2023) also showed significant correlation of modules in networks from more stressed communities to studied contaminants, with keystones (defined by higher connectivity) belonging to Rhodobacteraceae, *Nitrospira* and *Arenimonas*, as previously reported in other studies (e.g. Li et al. 2020). Members of these groups were also identified in this study as keystones, connectors or drivers in the moderately and heavily disturbed sites. This indicates a strength and validity of the approach taken here, consisting of using network topology for the identification of potential indicators of stress, across different types of sediment types and stress.

A higher proportion of negative *v* positive correlations with better EQ (lower stress) has also been reported previously, in agreement with our results (Hernandez et al. 2021). It has also been suggested that these negative correlations may contribute to a higher temporal community stability (Herren and McMahon 2018). In line with our results, lower network density with disturbance, has also been reported previously, in marine micro-, meio- and macrofauna impacted by offshore oil drilling (Laroche et al. 2018). Altogether, these results indicate that network topological properties can be useful as EQ indicators, although some studies also failed to observe any clear trends in this respect (Li et al. 2019; DiBattista et al. 2020). Standardised methods for modelling and comparisons of networks may, as suggested by Codello, Hose and Chariton (2022), could help to clarify if these differences in network topology are due to methodological or ecological differences between studies.

Keystones, connectors and drivers identified showed a wide taxonomic distribution across all CNs analysed, several of which also agreeing with previous studies. For example, we identified *Nitrospira*, *Novosphingobium* and Nitrosomonadaceae as drivers to good EQ, taxa that were previously identified as indicators of good status in freshwater stream sediments (Simonin et al. 2019) and negatively associated with heavy metal pollution (Li et al. 2020). Furthermore, *Thermodesulfovibrionia*, Anaerolineaecea members, Thiotrichaceae, clades SEEP-SRB1 and Spirochaeta 2, were identified as indicators of metal pollution in coastal sediments (Moreira et al. 2021), coinciding with the connectors and drivers identified here for Moderate and Poor CNs. Different members of the Desulfobacterota, meanwhile, were important players in all three CN.

Most Desulfobacterota undertake dissimilatory reduction of sulphate and other S compounds, driving the degradation of organic material during continuous or temporary anoxic or oxygen limited conditions, as well as contributing S cycling by sulphate removal and generation of hydrogen sulphide. As expected, they have therefore been identified as good indicators of organic enrichment and other types of disturbance in several studies (Keeley, Wood and Pochon 2018; Lanzén et al. 2020; Ye et al. 2023; Zvi-Kedem et al. 2024). However, oxygen limited conditions and local organic enrichment also occur naturally, explaining why high abundances of this phylum have also been reported in pristine tropical sites in Australia and India (Liao et al. 2020; Shah et al. 2021), and even in oxygen limited, carbon rich micro-habitats such as aggregates in pelagic waters (Siebers et al. 2024).

*Ca*. *Nitrosopumilus*, which has previously been identified as an indicator of good status (Lanzén *et al*. 2020, Moreira et al. 2021), was identified here as a driver from Good to Intermediate EQ, while *Draconibacterium,* previously identified as an indicator of poor EQ (Lanzén *et al*. 2020), was found to among the drivers to Good EQ here. These contrasting results highlight that driver taxa are not always applicable directly as indicators. Instead, limiting the identification of possible indicators to those that can be identified both as keystones and drivers, may be a more accurate approach, since it considers only taxa that are both important for CN structure and that may change its role and interaction structure if disturbance changes. Out of the nine such keystone-driver taxa identified here, six had been identified previously as indicators in the studied region and estuaries, corresponding to the expected EQ, namely Pseudomonadales-MBAE14 and Gammaproteobacteria-1013-28-CG33 for Good, Prolixibacteraceae for Moderate and *Acinetobacter*, Rhodocyclaceae and Macellibacteroides for Poor EQ (Lanzén et al. 2020).

PiCRUSt2 based prediction of the putative metabolic capabilities of the nine potential indicator (keystone-driver) taxa revealed five KOs with abundances corroborating those of their indicator host. These included the alanine-glyoxylate transaminase KO0830, mainly present in an environmental subgroup of Gammaproteobacteria in the sites with Good EQ, suggesting catabolism of alanine, which has also been predicted in pristine estuarine sediments from Far Northern Queensland (Shah et al. 2021). The alkaline phosphatase K01077, on the other hand, was mainly found in Poor EQ sites, in *Acinetobacter,* Pseudomonadales and Macellibacteroides. Although this enzyme is present across the tree of life and is a key enzyme in marine environments, regardless of P availability. Counter-intuitively, eutrophic habitats often show higher phosphatase activity, in spite of high concentrations of inorganic P (Hoppe HG 2003), in accordance with the higher presence at such sites observed here. In fact, phosphatase activity has even been suggested as a bioindicator of eutrophication (Taga and Kobori 1978) and shown to be relatively high compared to most marine habitats also in less nutrient-enriched, temperate, macrotidal estuaries similar to our Good EQ sites (Labry et al. 2016). (Costa, Carolino and Caçador 2007) have reported similar results, as well as higher activity in urbanized and industrial areas of estuaries.

In conclusion, this study illustrates how studying prokaryotic communities and modelling their ecological consensus association networks based on eDNA metabarcoding data from time series can provide additional insights into the environmental status of estuarine microbenthic communities. Time series also have the advantage of revealing temporal community stability, which may be impacted by disturbances, as observed here in one of the Moderate EQ sites. While less impacted sites showed higher stability, longer time series than those studied here would also provide a baseline for seasonality, as indicated for taxa like Thaumarchaeota and Desulfobacterota.

## Acknowledgements

We would like to thank Iñaki Mendibil for his assistance in the eDNA isolation, amplification and library preparation. This is contribution number XXXX of AZTI, Marine Research, Basque Research and Technology Alliance (BRTA).

