## Supplementary figures and images for "Contrasting dynamics and biotic association networks in estuarine microbenthic communities along an environmental disturbance gradient"

### Supplementary Figure 1

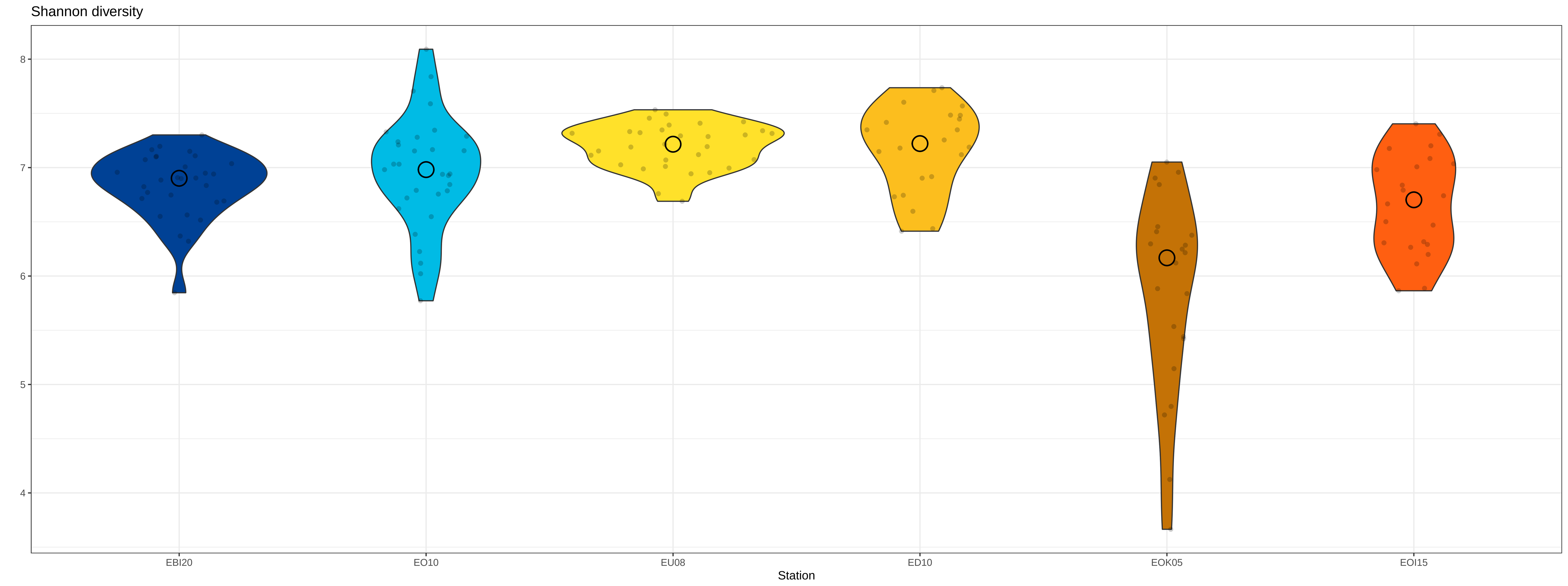

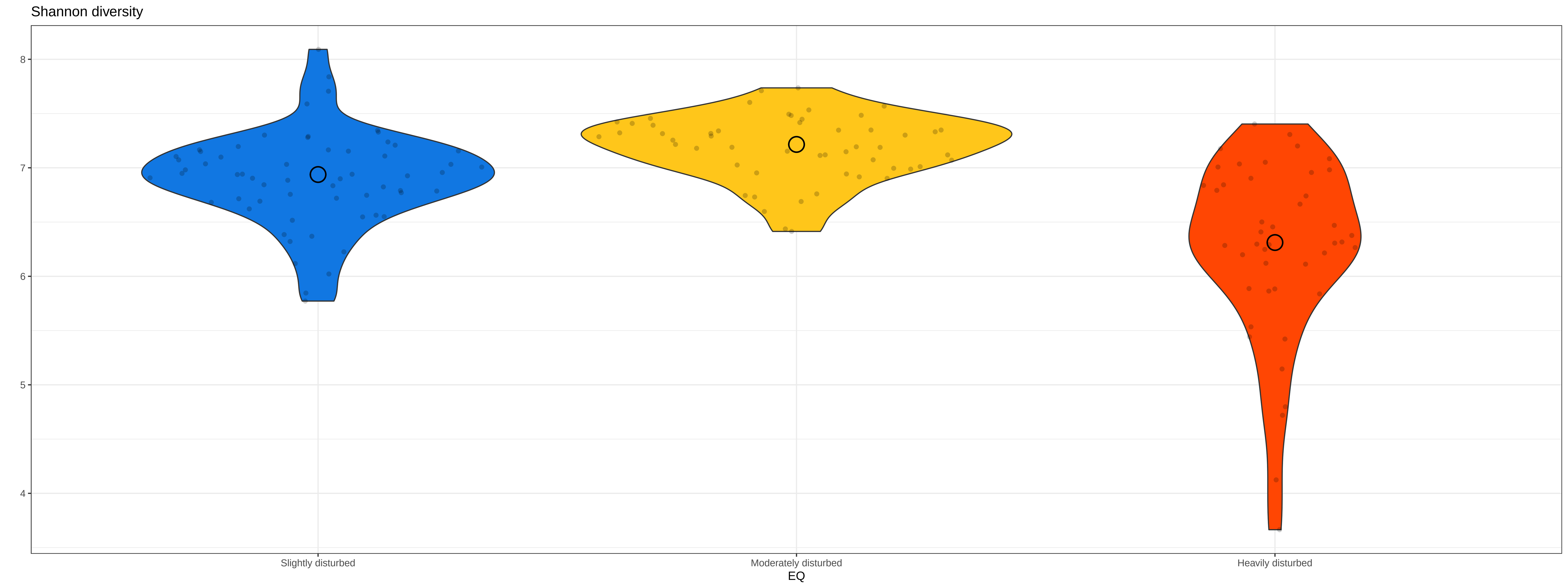

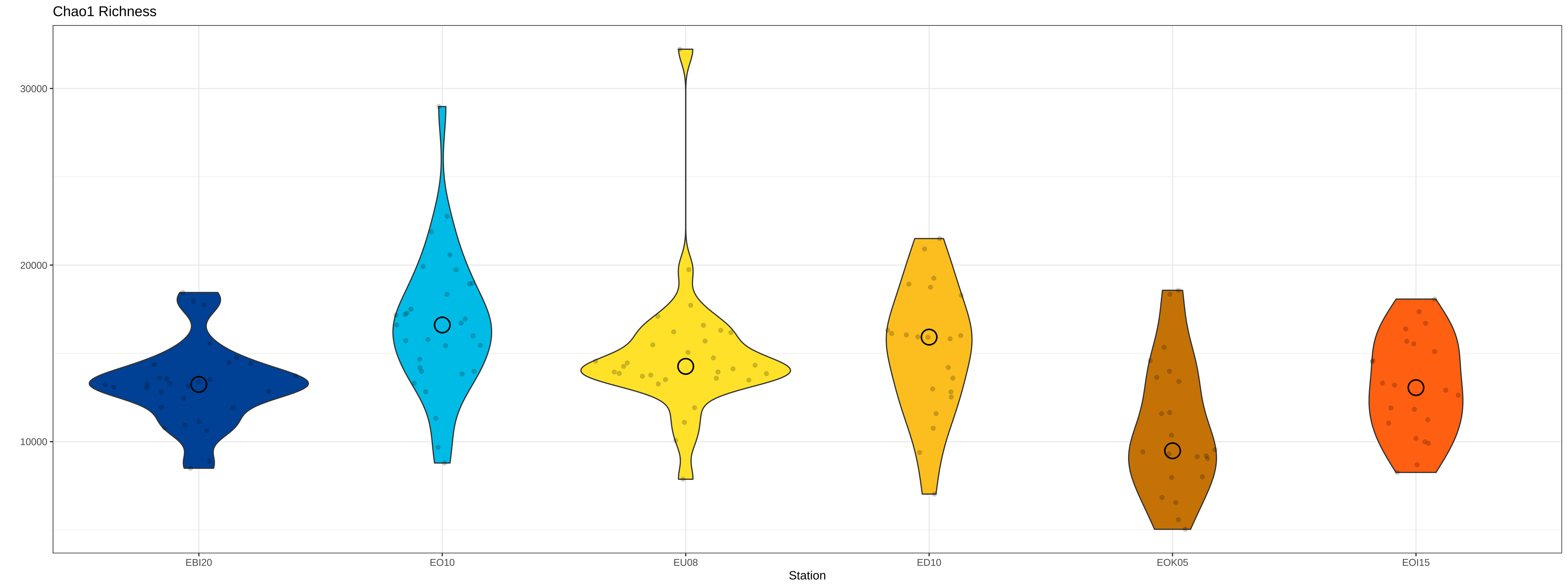

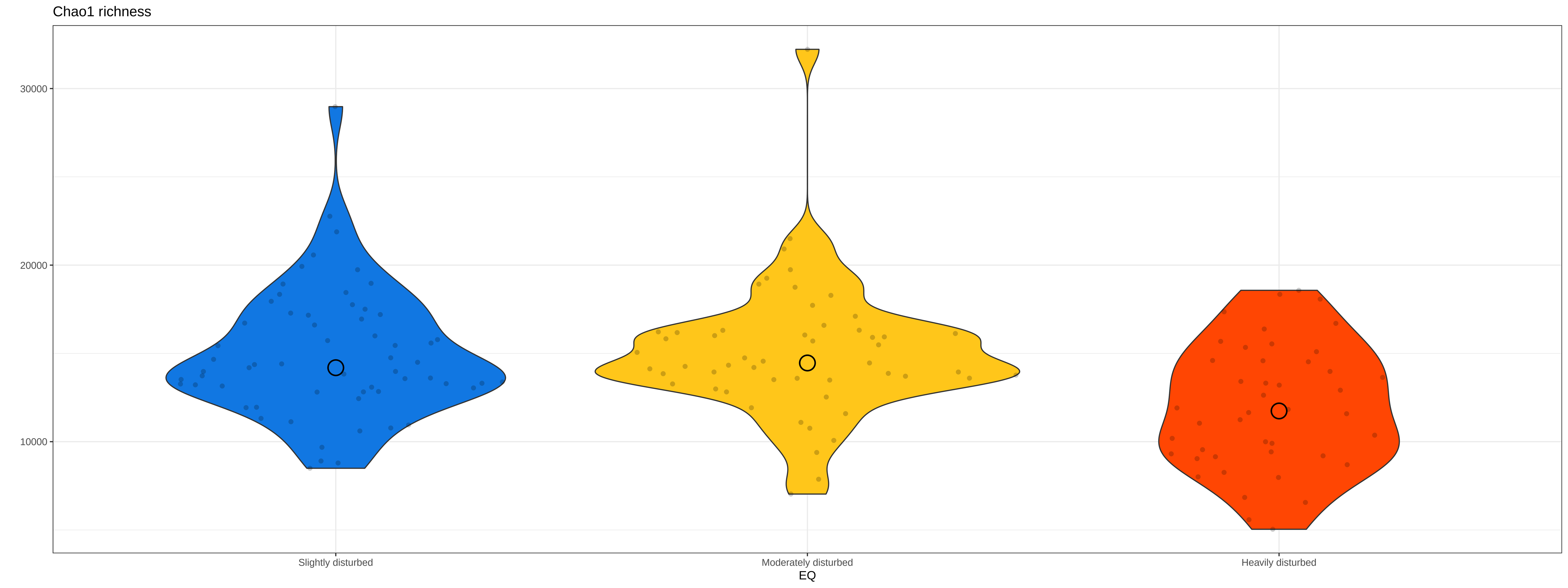

### Supplementary Figure 2

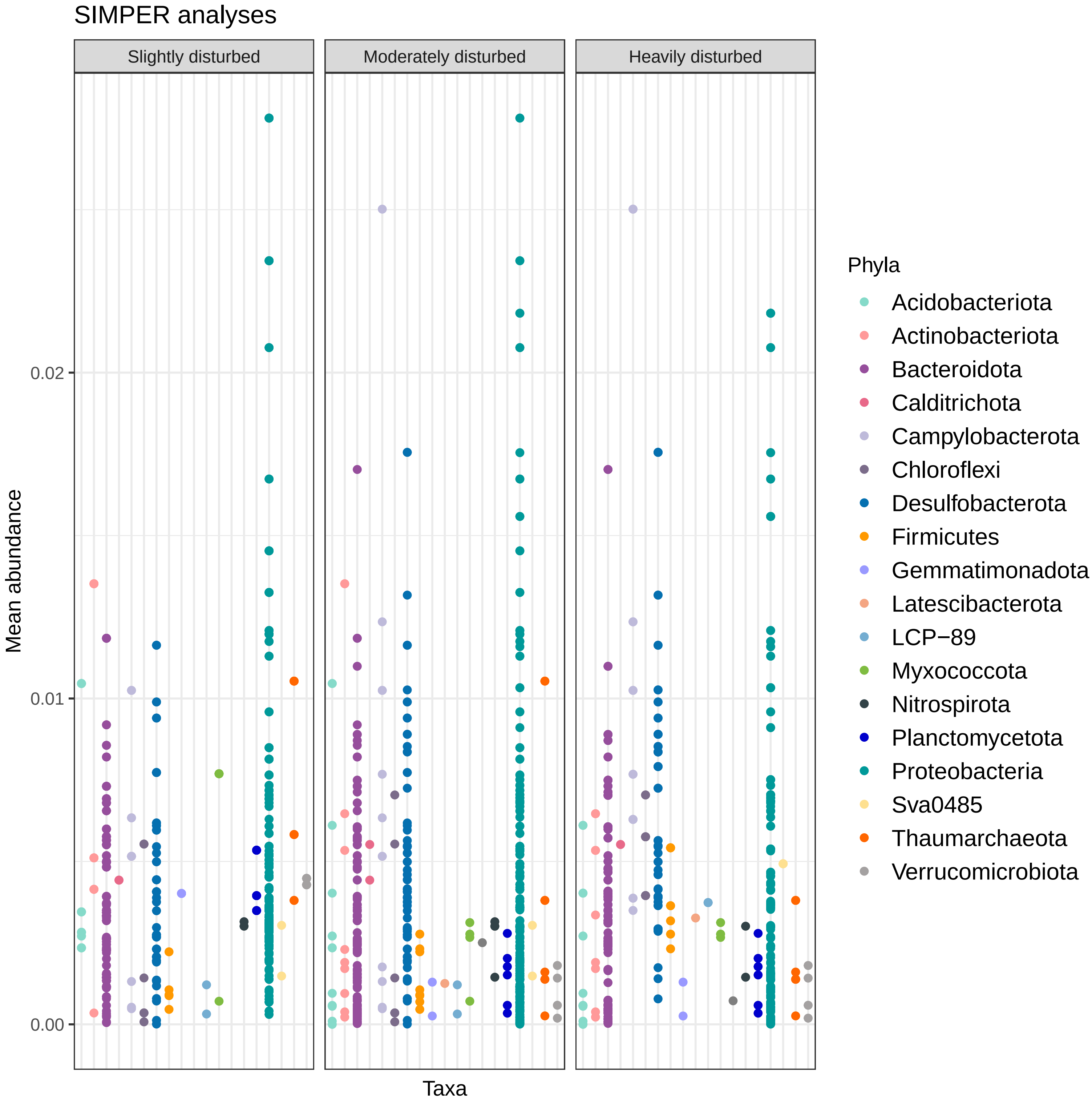
